# The effect of colonic pH on microbial activity and metabolite production using common prebiotics as substrates: an *in vitro* study

**DOI:** 10.1101/2023.03.29.534806

**Authors:** Zhuqing Xie, Weiwei He, Alex Gobbi, Hanne Christine Bertram, Dennis Sandris Nielsen

## Abstract

The interplay between gut microbiota (GM) and host via degradation of dietary components leading to the production of metabolites such as short-chain fatty acids (SCFAs) is affected by a range of factors including colonic pH and carbohydrate source. Here we investigate how differences in colonic pH influence GM composition and metabolite production using different substrates including inulin, lactose, Galactooligosaccharides (GOS), and Fructooligosaccharide (FOS) in an *in vitro* colon setup. We investigated 3 different pH regimes (low, 5.2 increasing to 6.4; medium, 5.6 increasing to 6.8 and high, 6.0 increasing to 7.2) and found that *Bacteroides* spp. decreased but *Bifidobacterium* spp. became abundant under low pH regimes, suggesting complex interactions of the bacterial community in the face of pH fluctuations in the colon. The butyrate producers *Butyricimonas* and *Christensenella* were enriched at higher pH conditions, where also butyrate production was increased using inulin as substrate. The relative abundance of *Phascolarctobacterium*, *Bacteroides*, and *Rikenellaceae* was also increased at higher colonic pH, which was accompanied by increased production of propionate using GOS and FOS as substrate. The gastrointestinal factors are linked in a complex network, where microbial activity leads to the production of SCFAs and other compounds that influence pH, which in turn seems to influence microbial activity. Taken together, our results show that dynamic changes in colonic pH under *in vitro* simulated conditions have a strong effect on gut microbial activity with SCFA production being higher at colonic pH conditions close to neutral.

## Introduction

The human colon harbors a complex microbial community that influences host physiology and metabolism. The gut microbiota (GM) can utilize different otherwise indigestible dietary fibers through fermentation thereby generating a variety of metabolites including short-chain fatty acids (SCFAs), succinate, lactate, methane, and hydrogen.^1^ Acetate, propionate, and butyrate are the major SCFAs and are of particular interest not only as an energy source, but also via exhibiting benefits to the host including enhancing satiety, suppressing appetite, and alleviating inflammation.^2^ The interplay between GM and the host via the degradation of fibers to produce SCFAs is highly complicated and can be affected by several factors, including colonic pH and carbohydrate source.^3, 4^ Colonic pH differs from person to person as well as between different colonic segments, where the pH of the proximal (5.4-5.9) and transverse (6.1-6.4) colon is lower than the distal colon which is close to neutrality (6.4-8.0).^5, 6^ Abundant bacteria in the proximal colon favor saccharolytic fermentation leading to SCFA production which decreases the colonic pH.^7^ In contrast, proteolytic fermentation dominates the distal colon where the available carbohydrates are depleted leading to an increase in pH.^8^ This might influence host health and physiology, as cations (mainly Ca^2+^) have higher bioavailability at lower pH^9^ and with low colonic pH generally protecting against microbial pathogens.^10^

Changes in pH influence not only the bacterial community^11^ but also metabolite production,^4^ which further connects with colonic function.^12, 13^ It has been found that acidic pH overall supports the growth of bacteria belonging to *Firmicutes* phylum whereas many *Bacteroides* members are more sensitive to acidity.^14^ *Escherichia coli*, a potentially pathogenic species in the human colon, can be inhibited by a decreased pH under *in vitro* colonic environments, whereas bifidobacteria and lactobacilli are favored by more acidic conditions.^14, 15^ Furthermore, colonic pH has been reported to influence the production of specific metabolites with butyrogenic reactions being favored at a slightly acidic pH, which is in contrast with propionate production that often occurs at a neutral pH.^16^ Acetate on the other hand can readily be produced at different pHs via the activity of various acetate-producing microbial species having different pH optima.^3, 17^ As a precursor for the production of other SCFAs, especially propionate and butyrate, lactate usually does not accumulate in the colon to any larger extent, and its concentration is closely related to the abundance of lactate-utilizing bacteria. However, some of the lactate utilizers are rather sensitive to acidic pH which may result in lactate accumulation under moderately acidic environments.^18^

The carbohydrate source is another factor that influences metabolite production and GM response. Reichardt et al. performed *in vitro* simulated colon batch fermentations of 15 different dietary fibers including glucans, pectins, hemicellulose, and fructans at two starting pH values (5.5 and 6.5), and found that butyrate was produced at the lower pH (5.5) for most substrates, which was in contrast with propionate production that in general was impaired at the lower pH.^16^ Chung et al. investigated SCFA production and microbial community response to inulin or pectin as substrate in pH-controlled fermentors inoculated with fecal matter and found that reducing pH from 6.9 to 5.5 promoted the abundance of *Faecalibacterium prausnitzii* replacing otherwise dominating *Bacteroides* spp.^4^ Ilhan et al. used glucose, fructose, or cellobiose as the single carbon sources in batch fermentors using fecal slurry as inoculum under three different starting pHs (6.0, 6.5, and 6.9).^19^ It was found that pH had substantial impact on lactate utilizers and producers, which was accompanied by lactate accumulation at pH 6.0. Besides, microbial diversity was driven not only by pH but also by carbon source, as cellobiose generated more acidity and had a pronounced effect on GM structure when compared with other substrates. These findings suggest that in addition to substrate type, gut environment, especially pH, can also strongly influence the metabolites being produced and the interspecies competition in the gut.

Inulin, fructooligosaccharides (FOS), and galactooligosaccharides (GOS) are typical prebiotics that have been found to promote especially *Bifidobacterium* spp. but also different lactobacilli in the gut.^20^ Despite the findings mentioned above, there is still only limited knowledge on how colonic pH may influence microbial community and metabolite production in the gastrointestinal tract. Here, we used the CoMiniGut *in vitro* colon model to investigate GM composition and SCFA production as influenced by pH after 24 h of *in vitro* simulated colon passage with the common prebiotics inulin, FOS, GOS, and lactose as substrates. Fresh stool samples from three donors with different fecal pH values (6.4, 6.8, and 7.2 respectively) were collected, and three dynamic pH programs (proximal to the distal colon) during 24 h of fermentation were designed which represented low (5.2-6.4), medium (5.6-6.8) and high (6.0-7.2) colonic pH individuals. Metabolite outcomes after 24 h of fermentation for each fecal slurry under the different pH regimes were examined by ^1^H NMR metabolomics, and GM shifts were traced using 16S rRNA gene amplicon sequencing.

## Results

### Colonic pH significantly influenced *in vitro* simulated GM structure, but the influence of fecal donor and substrate was more pronounced

After 24 h of *in vitro* simulated colon passage the number of Observed species as well as the Shannon diversity index increased with colonic pH for all substrate types (inulin, lactose, GOS, and FOS) (see Figure 2A and 2B for average across donors and Figure S1 for individual donors). Bray-Curtis dissimilarity-based db-RDA analysis (Figure 2C) showed clear groupings of the *in vitro* simulated colon fermentation samples according to pH, which was confirmed by pairwise PERMANOVA (Figure 2E and Figure S2 C-D). However, pH explained only 3.48% of the total variance (Figure 2D) relative to the effect of the donor (35.99%) and substrate (8.34%). In accordance with this, PCoA plots based on both Bray-Curtis dissimilarity and Jaccard similarity metrics showed strong donor effects (Figure S2 A-B) which is also reflected by the actual microbiome composition as seen in Figure 3. Overall, all four tested substrates led to a higher relative abundance of bifidobacteria relative to inoculums. Low *in vitro* simulated colonic pH especially promoted the proliferation of bifidobacteria when inulin was used as substrate (Figure S3).

**Fig. 1:**
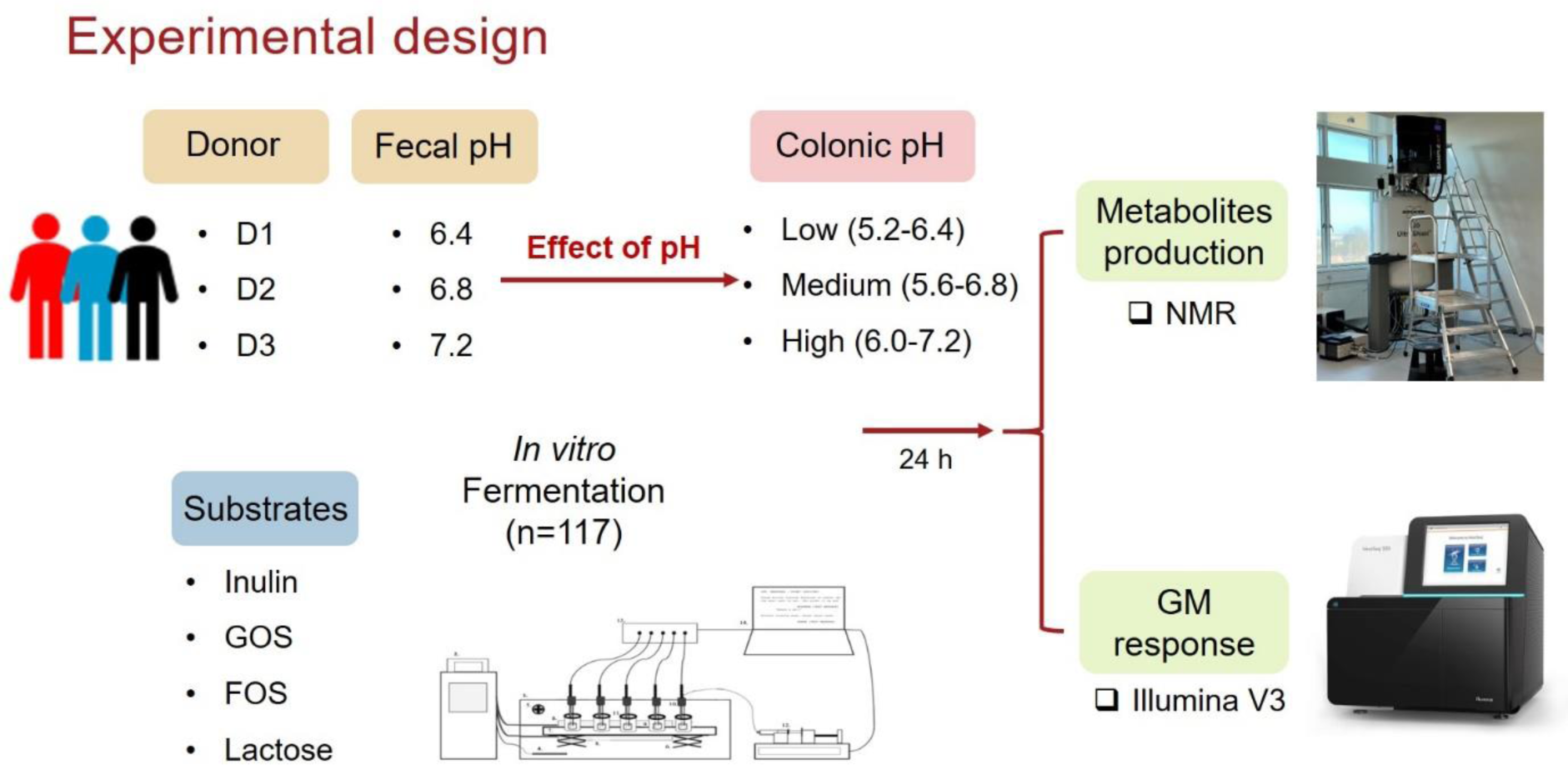
Diagram illustrating the experimental design. Fresh stool samples from three healthy donors with different fecal pH values (6.4, 6.8, and 7.2, respectively) were collected, and three dynamic pH programs during 24 h of fermentation were designed which represent the low (5.2-6.4), medium (5.6-6.8) and high (6.0-6.8) pH changes from the proximal to the distal colon. Single substrates (inulin, lactose, GOS, or FOS) were added to each chamber for colonic fermentation. Metabolite outcomes after 24 h of fermentation for each fecal slurry under the corresponding pH and abnormal colonic pHs were detected by ^1^H NMR, and GM shifts were traced using the V3 region of the 16S rRNA gene amplicon sequencing by Illumina, NextSeq.

**Fig. 2.**
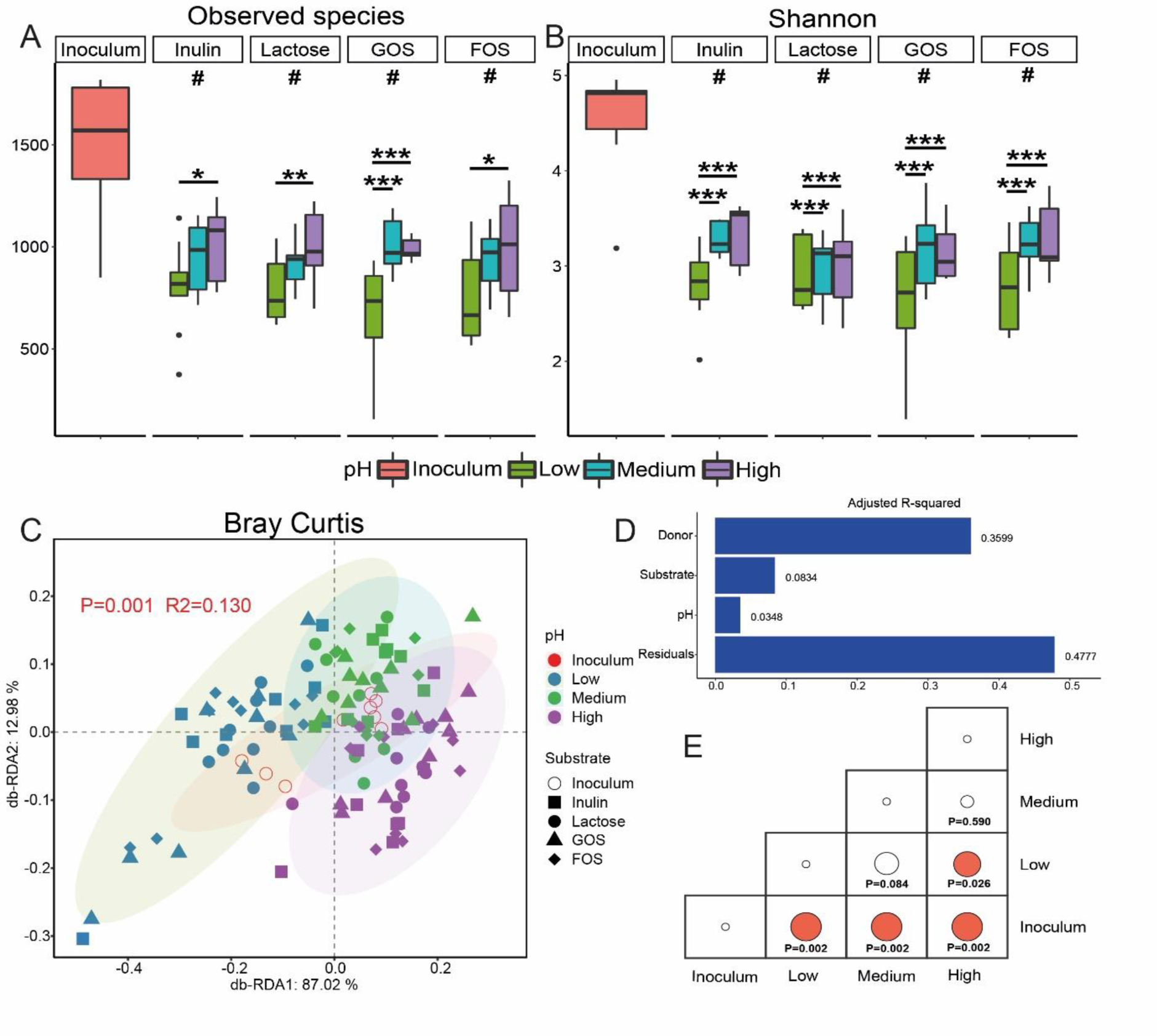
The influence of colonic pH on GM diversity after 24 h of fermentation. Observed species (A) and Shannon index (B) of the microbial community, Db-RDA biplot (C) showing microbial variance explained by colonic pH with adjusted R2 (D) and pairwise PERMANOVA tests (E) on Bray Curtis metrics. A t-test was applied to determine the influence of pH on alpha diversity for three inocula with a single substrate. Significant differences between changed colonic pH are labeled with * (p < 0.05), ** (p < 0.01) and *** (p < 0.001), respectively. Significant differences relative to inoculum are labeled with # (p < 0.05).

**Fig. 3.**
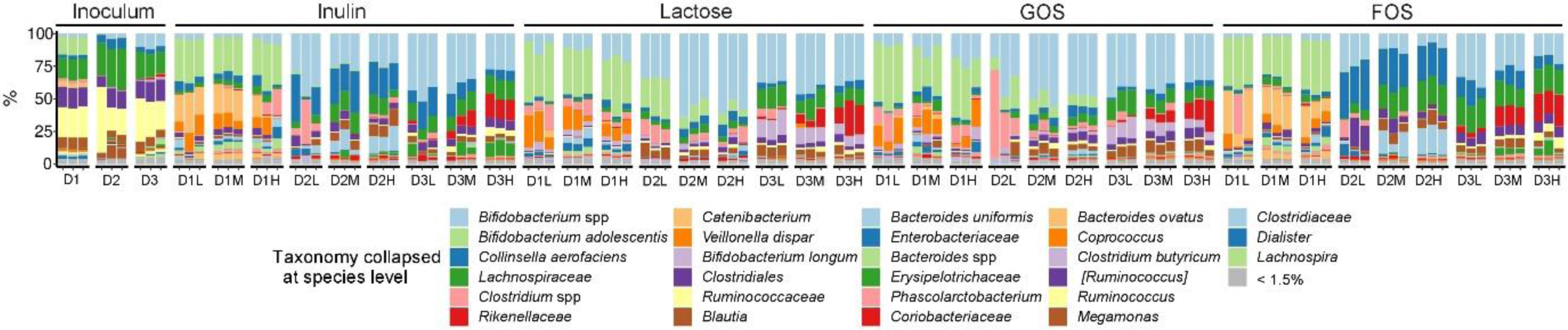
Summarized species-level gut microbiota composition of individual fermentation samples with different substrates.

### Low *in vitro* simulated colonic pH strongly influenced microbiome composition

Deseq2 was carried out to determine differences in the *in vitro* simulated GMs under different colonic pH regimes (Figure 4). When comparing the effect of pH, it is evident that low pH strongly influences the microbiome, while only a *Mogibacteriaceae* member was found to differ between the medium and the high pH regime. In summary, acidic pH promoted most *Firmicutes* members (*Clostridium* members, *Lutispora*, and *Dialister*) but suppressed phylum *Bacteroidetes* members including *Bacteroides*, *Butyricimonas*, and *Rikenellaceae* relative to high pH conditions. Low pH conditions lead to the decreased relative abundance of *Christensenellaceae*, *Phascolarctobacterium, Holdemania*, *[Mogibacteriaceae]*, and *Christensenella* for all four tested substrates relative to the medium and high pH regimes.

**Fig. 4.**
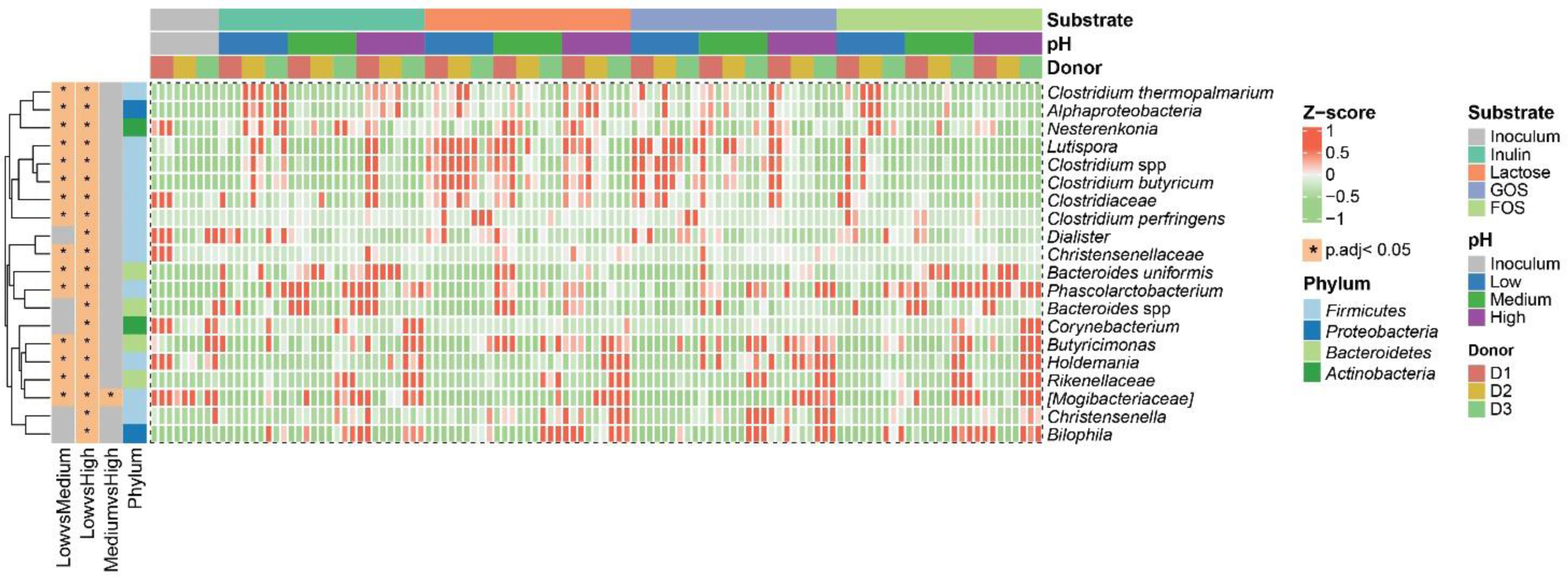
Differentially abundant taxa between changed colonic pH. The top 20 differentially abundant taxa with the lowest p-adjust value were selected by Deseq2 and labeled with * in the heatmap, indicating the significantly altered taxa from the respective between-group comparison. Row-wise Z-score scaling was conducted in the heatmap visualization, showing the normalized relative abundance by the mean of the specific taxa across all samples.

### Higher *in vitro* simulated colonic pH promoted the production of SCFAs in a donor- and substrate-dependent manner

Higher *in vitro* simulated colonic pH favored SCFA production, especially when grown with inulin and FOS as substrate (*p* < 0.05) (Figure 5). Inulin particularly promoted butyrate formation, while FOS and GOS promoted propionate formation (Figure 5), which was confirmed by the OPLS-DA model and S-line plots of NMR spectra (Table S1 and Figure S4). Acetate production appeared to be less influenced by pH, as a pH effect only was observed for inulin as substrate with a significant promotion in acetate concentration at the medium pH. Formate and lactate, two intermediate products/substrates in SCFAs formation, were only detected in relatively low amounts here, and lactate accumulation during GOS-based fermentation was reversed by increased colonic pH (Figure 5F). Again, donor-specific differences were observed with respect to both total SCFA formation as well as specific SCFAs like acetate and propionate being promoted by higher pH with GOS as the substrate for donors 2 and 3, whereas SCFA production by donor 1 was not affected by pH to any larger extent when grown on GOS (Figure S5). Besides, lactose presented apparent individualized SCFA production, where the corresponding pH regime for Donor 1 (low fecal pH) and Donor 3 (high fecal pH) favored higher acetate and propionate production, respectively.

**Fig. 5:**
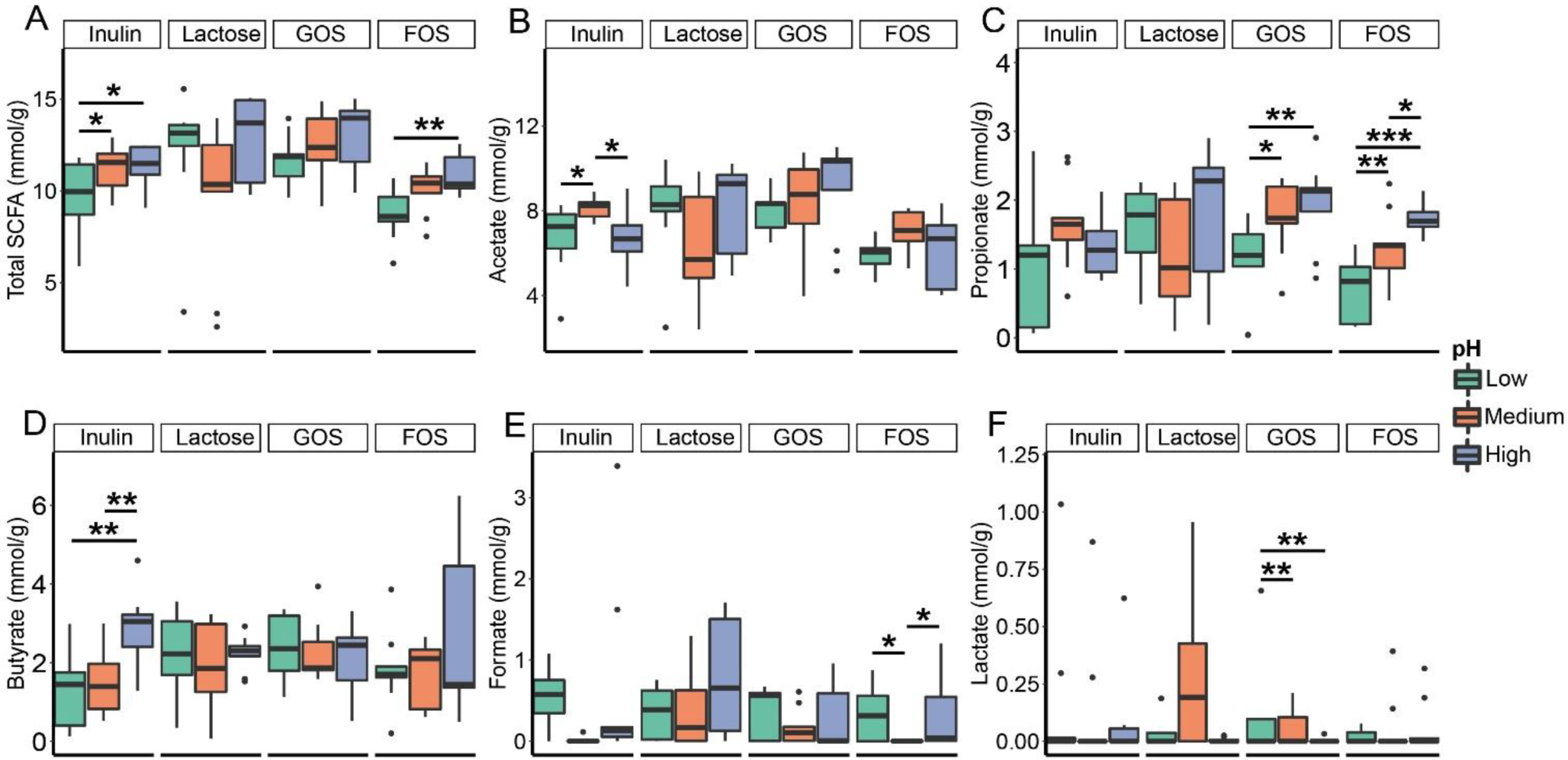
The influence of colonic pH on metabolite production after 24 h of fermentation. A-F represents total SCFA, acetate, propionate, butyrate, formate, and lactate production, respectively. A t-test was applied to determine the influence of pH on metabolite production with a single substrate. Significant differences between changed colonic pH are labelled with * (p < 0.05), ** (p < 0.01) and *** (p < 0.001), respectively.

### Propionate and butyrate production is associated with specific bacterial taxa and pH-levels

Associations between specific bacterial species and metabolite production were determined by Pearson’s correlation analysis and 138 significant pairs were found (Table S2). As seen from Figure 6A, the high relative abundance of *[Mogibacteriaceae]*, *Phascolarctobacterium*, *Rikenellaceae*, and *Bacteroides* spp. at the high colonic pH was positively correlated with propionate production, while *Butyricimonas*, which also showed higher relative abundance with higher pH, presented a positive relationship with butyrate concentration. Besides, co-occurrence analysis (Figure 6B) was conducted to explore the interactions among species where the relative abundance was influenced by simulated colonic pH. Further, several abundant microbial members with strong positive co-occurrence (e.g. *Lachnospiraceae*, *Ruminococcus*, *Clostridiales*, *Coprococcus*, and *Blautia*) consistently correlated with more propionate/butyrate production (Figure 6A).

**Fig. 6:**
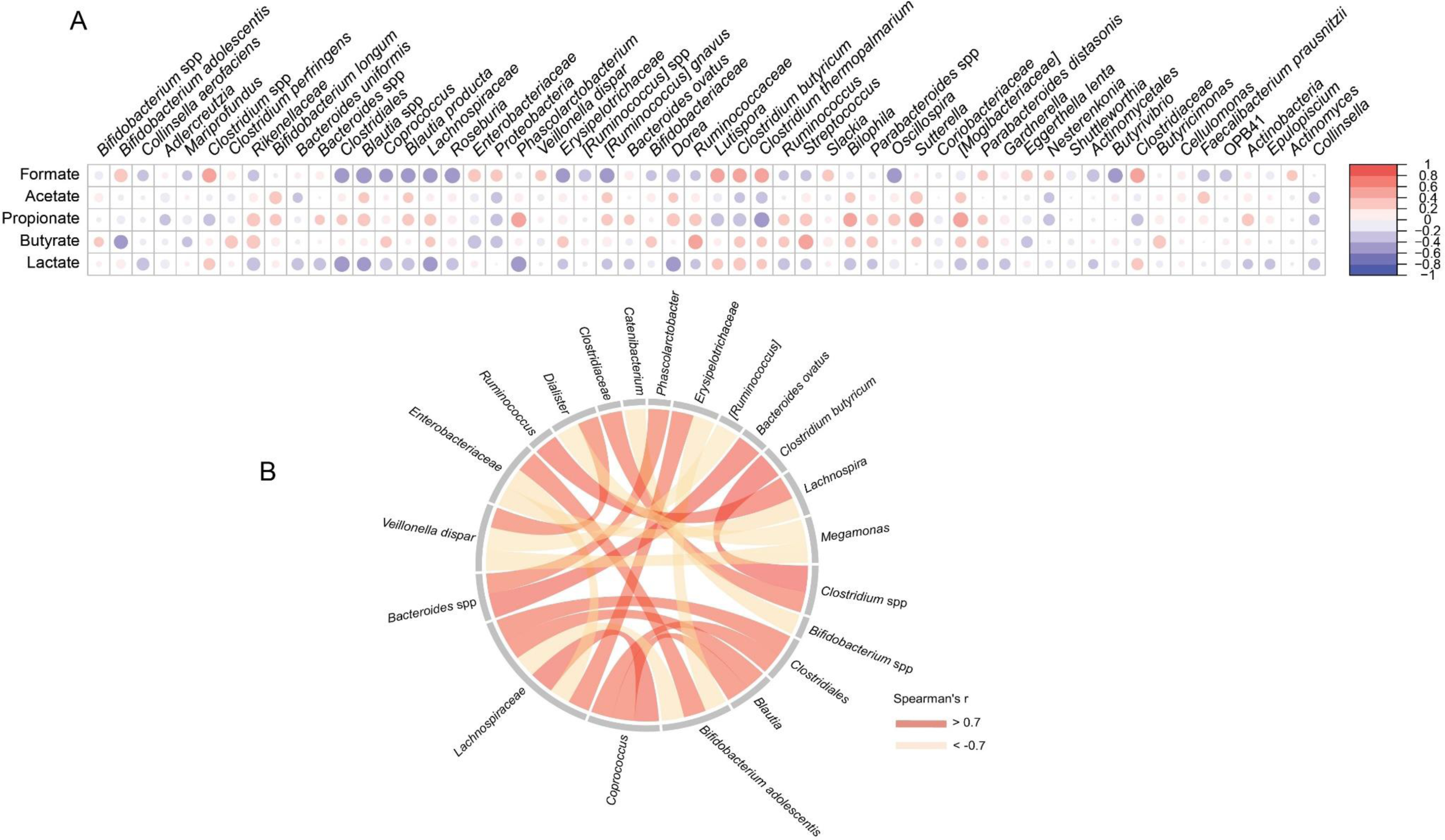
Correlation between microbial taxa relative abundance and metabolites (formate, acetate, propionate, butyrate, and lactate) concentrations (A) and inter-species co-occurrence (B). The minimal prevalence (among the sequenced samples) of one given microbial taxa was set to 30%. Species with a mean relative abundance >1.5% were chosen for co-occurrence analysis, and Spearman’s rank correlation coefficient of > 0.7 is shown in the chord diagram.

## Discussion

Indigestible dietary compounds such as fibers and other complex carbohydrates benefit host physiology via the production of key bacterial metabolites like SCFAs produced in the colon,^21^ which in turn also influence colonic pH.^22^ Wide inter-individual differences in colonic pH (5.3-8.0) and gut transit time have been reported,^6^ but it still remains poorly understood whether factors such as the dynamic pH increase from the proximal to distal colon could influence interspecies bacterial balance and metabolites formation. To investigate this, we employed an *in vitro* colonic model with dynamic pH control simulating the pH conditions of the ascending, transverse, and descending colon. As inoculum, we used fresh stool samples from three healthy adults representing different fecal pH-levels (6.4, 6.8, and 7.2) and thus also representing individuals with low, intermediate, and high colonic pH, respectively.

Interestingly, the alpha diversity index showed the same trend for inulin, lactose, GOS, and FOS substrate groups, as the Observed species and Shannon Diversity index were higher at a higher colonic pH. The high alpha diversity index observed here is consistent with previous results^23^ and is generally anticipated to be associated with resisting microbial perturbation and diverse functions in the gut.^24, 25^ The significant reduction in alpha diversity (Observed species and Shannon index) of four substrates relative to inoculum (Figure 2A-B) might result from adding a single carbon source in the medium under simplified *in vitro* simulations, which contrasts with the human diet containing complex components.^26^ In our study, colonic pH significantly influenced the microbiota composition, but to a smaller degree relative to the effect of donor and substrate (Figure 2D). It is well established that specific substrates could boost the population of specific bacteria but the microbial patterns showed a clear inter-individual variability.^16^ The combination of complex factors determined the long-term stability of individual GM according to a previous report,^27^ likely explaining the importance of the donor effect on the GM structure (Figure 2D) as well as the individualized-microbial composition (Figure 3) in our results. Our findings showed that substrate has a great impact on bacterial community relative to the pH, which is inconsistent with a previous report,^19^ and the explanation might be found from the complexity of the individual bacterial composition and interactions with different diet components *in vivo*.

The high relative abundance of *Actinobacteria* members such as *Bifidobacterium adolescentis* and other bifidobacteria as well as *Collinsella aerofaciens* after fermentation using inulin, lactose, GOS, or FOS (Figure 3) as substrate confirmed the bifidogenic effects of these common prebiotics.^28, 29^ For example, *Collinsella aerofaciens* is well-known for fermenting a variety of carbohydrates and producing SCFAs in the human colon.^30^ Besides, we observed that the high abundance of *Bifidobacterium* spp. at low pH was associated with inulin and FOS substrates for donor D2 and D3 (Figure 3 and Figure S3), indicating that the proliferation of *Bifidobacterium* spp. was closely related to colonic pH. This is in agreement with previous reports showing higher *Bifidobacterium* spp. relative abundance at pH 5.5 compared to pH 6.5 with arabinoxylan^16^ and inulin^4^ as a single carbohydrate source. This might be explained by the enhanced activity of *Bifidobacterium* spp. enzymes at acidic pH conditions.^11^ It can be speculated that the bifidogenic effect of substrates like inulin observed *in vivo*^31^ might result from the synergistic effect of such substrates leading to increased SCFA production that lowers colonic pH, which again renders *Bifidobacterium* metabolism more efficient.

Also the production of SCFA differed in a donor and substrate-dependent manner, but less pronounced than what was observed for the GM profiles (Figures 3, 4, 5, and S5). Similarly, Reichardt et al. (2018) also found that the GM profile was more donor-dependent than the SCFA profile.^16^ This is the result of a large number of colonic bacteria capable of utilizing substrates and producing acetate, propionate, and butyrate as three main fermentation products, thus leading to functional redundancy with respect to metabolite production relative to the GM composition. For inulin, GOS, and FOS the SCFA production increased with higher simulated colonic pH. This is in support of previous findings^16^ that especially acetate and propionate production is higher at pH 6.5 relative to pH 5.5. The increased butyrate production from inulin at the higher simulated colonic pH-levels tested positively correlated with the relative abundance of the butyrate-producers *Butyricimonas* (Figure 6A)^32^ and *Christensenella* (Figure 4),^33, 34^ and consistent with previous findings, we observed positive correlations between butyrate production and the abundance of *Coprococcus*,^35^ *Lachnospiraceae*,^36^ and *Ruminococcus*^37^ members, which have previously been reported as potentially butyrogenic bacteria. Propionate production and *Phascolarctobacterium*, *Bacteroides*, and *Rikenellaceae* relative abundance were linked and both enhanced at the higher colonic pH levels tested for the GOS and FOS groups. *Rikenellaceae* members are prominent fiber fermenters in the human colon resulting in the production of propionate.^38^ *Phascolarctobacterium* spp. are succinate-metabolizing bacteria co-existing with *Bacteroides* members in the gut and producing substantial amounts of acetate and/or propionate which is in accordance with the strong association between these factors observed in Figure 6B.^14, 16, 39^ Previously, decreased GM alpha diversity^4^ has been observed when pH gets below pH 6.0, which apparently is linked to reduced growth of *Bacteroidales* at relatively low colonic pH, possibly indicating that the effect of colonic pH on GM could be related to alterations within the *Bacteroidales* community. In line with this, lactate accumulation in GOS-based fermentations at low simulated colonic pH matched with a corresponding reduction of propionate levels, which can be explained by the inter-species competition between lactate-producing *Bifidobacterium* and –utilizing bacteria *Bacteroides*.^17, 18^

## Conclusion

In summary, we found that the influence of colonic pH on SCFA production is linked to concurrent changes in the bacterial community profile in a donor and substrate-dependent manner. Our results strongly indicate that colonic substrates such as dietary fibres influence GM composition and metabolite production, not only by being selectively utilized by specific microbes, but also because of their SCFA production, which in turn also influence colonic pH and overall GM composition and activity.

## Materials and methods Materials

FOS (F8052) was purchased from Sigma-Aldrich Chemical Co. (St. Louis, MO, USA). Inulin (YI012742001) and GOS (OG321341901) were purchased from Biosynth Carbosynth (Berkshire, UK). Lactose (VM922157008) was purchased from Merck KGaA (Darmstadt, Germany). All reagents used in phosphate-buffered saline (PBS) and basal colon media (BCM) ^40^ were of analytical grade.

### Fecal samples collection and pH determination

Three donors with different fecal pHs (6.4, 6.8, and 7.2) were selected for this study to provide fresh stool samples for the fermentations. The donors were all healthy adults without dietary restrictions and antibiotic treatments over the past three months (Ethical Committee of the Capital Region of Denmark registration number H-20028549). The fecal samples were handled under anaerobic conditions in an anaerobic chamber (AALC model, Coy Laboratory Product, atmosphere ∼ 93% (v/v) N_2_, 2% H_2_, 5% CO_2_). Individual stool samples were transferred to the chamber within minutes after collection and homogenized with sterilized PBS/20% glycerol (v/v) at a ratio of 1:1 for 120 s using the Stomacher (Stomacher 400; Seward, Worthing, UK). The glycerol stocks were aliquoted into cryo-vials and stored at −60 ℃ before use as fecal inoculum for the fermentations. One part of the individual feces was mixed with distilled water at a ratio of 1:9 after collection, and the pH of the mixture was determined using a pH meter calibrated on the same day (FC240B, Hanna Instruments, UK).

### *In vitro* simulated colonic fermentations

The fermentations were performed using the previously described CoMiniGut *in vitro* colon model with minor modifications to the procedure.^40, 41^ The experimental design is summarized in Figure 1. All tested fermentation conditions were conducted in triplicates. Briefly, fecal glycerol stocks from three donors were thawed and further diluted with sterilized PBS in a ratio of 1:4. All fermentations were carried out in 4.5 mL BCM containing 50 mg substrate (inulin, FOS, GOS, or lactose) and 0.5 mL fecal slurry. Anaerobic conditions during fermentation were achieved by positioning an anaerogen compact sachet (AN0010W; ThermoScientific, Waltham, MA, USA) in the reaction chamber. Resazurin-soaked indicators (Anaerobe Indicator Test; Sigma-Aldrich, St. Louis, MO, USA) were used to signal anaerobiosis. Three dynamic pH programs (low, 5.2 increasing to 6.4; medium, 5.6 increasing to 6.8 and high, 6.0 increasing to 7.2) simulating the pH from the proximal colon to the distal colon were controlled by connecting the pH meter with a laptop running in Matlab scripts which regulates a multichannel syringe pump charged with syringes adjusting pH by adding 2.5 µL aliquots of 1 mM of NaOH pr. bolus. After 24 h of fermentation, fermenta were collected for further metabolite determination and GM analysis.

### ^1^H NMR spectroscopic analysis and metabolite quantification

^1^H NMR spectroscopy of collected fermenta was performed using Bruker Avance IVDr NMR spectrometer (Bruker BioSpin, Rheinstetten, Germany) equipped with a 5 mm ^1^H-optimized double resonance broad-band probe and operating at a frequency of 600.13 MHz for ^1^H. Samples for ^1^H NMR spectroscopy were prepared according to a procedure described by He et.al^42^ with minor modifications. In brief, 500 µL of fermentation sample was vortexed and filtered by centrifugation at 4 ℃, 14 000 g for 30 minutes using Amicon Ultrafilters (Merck Millipore Ltd., Cork, Ireland), and 300 µL supernatant was mixed with 300 µL phosphate buffer in D_2_O (pH 7.4) containing trimethylsilylpropanoic acid (TSP) (0.01% w/w) in 5 mm NMR tubes. ^1^H NMR spectra were obtained at a temperature of 300 K using a one-dimensional NOESY pulse sequence with following acquisition parameters: relaxation delay: 4s, spectral width: 7212 Hz, data points: 32k, and number of scans: 32. The free induction decays with a line-broadening factor (0.3 Hz) were adopted before Fourier transformation. Phase adjustments and baseline correction of obtained ^1^H NMR spectra were performed in the Topspin 3.6.2 software. Multivariate data analysis (MVDA) including principal component analysis (PCA) and supervised orthogonal projections to latent structures discriminant analyses (OPLS-DA) were conducted on binned ^1^H NMR spectra (bin width = 0.005 ppm) after removal of the residual water signal (4.75-7.90 ppm). The S-line plots of the OPLS-DA models were employed to examine the spectral differences between the low and high pH programs of the total samples and fermenta with inulin as a carbohydrate source, respectively. In addition, identification and quantification of metabolites from the ^1^H NMR spectra obtained for the 24 h fermenta were employed by Chenomx (Version 8.6, Chenomx Inc., Alberta, Canada). For metabolite production including the total SCFA, a t-test was applied for pairwise comparison of different pH programs in each substrate after 24 h of fermentation.

### DNA extraction, library preparation, and Illumina sequencing

The microbiome composition of *in vitro* simulated colonic fermentations as well as the fecal inoculums were characterized by 16S rRNA gene amplicon sequencing. DNA extraction was performed using the Micro Bead beat AX kit (A&A Biotechnology, Poland) following the manufacturer’s instructions. Qubit dsDNA BR Assay Kit (Thermo Fisher Scientific Inc., Waltham, USA) and NanoDrop ND-1000 Spectrophotometer (NanoDrop Technologies Inc., Wilmington, USA) were used for determining the concentration and purity of the extracted DNA. The V3 hypervariable region of the 16S rRNA gene was amplified using primers compatible with the Nextera Index Kit (Illumina, San Diego, CA, USA), and library preparation was performed according to the method published elsewhere.^43^ The PCR2 products were purified using AMPure XP beads (Beckman Coulter Genomic, CA, USA) and pooled in equimolar concentrations for sequencing using the Illumina NxtSeq platform.

### Bioinformatics and statistics

The raw Illumina data set containing pair-ended reads with matching quality scores were merged and trimmed in the USEARCH pipeline^43^ using fastq_mergepairs and fastq_filter scripts. UNOISE3^44^ was used to build zero radius Operational Taxonomic Units (zOTUs), and the Greengenes (13.8) 16S rRNA gene collection^45^ was used as a reference database for taxonomy assignment. Statistical analysis and plot visualization were performed by R (v 4.1.3). The feature table, taxonomy, metadata, and tree file were imported through phyloseq in R.^46^ For diversity analysis, samples were rarefied to 70 000 reads with the function “rarefy_even_depth” in phyloseq. A t-test was conducted to determine the statistical differences in alpha diversity (Observed species and Shannon index) between different pH programs of the specific substrates, and PERMANOVA was employed for determining GM structural changes based on Bray-Curtis dissimilarity and Jaccard similarity matrixes, respectively. Distance-based redundancy analysis (db-RDA) based on Bray-Curtis metrics was performed to discern the variance explained by colonic pH, and the effect size of other factors such as donor, substrate, and pH was tested by the function “adonis”. For GM composition, taxa were agglomerated at the species level, and the collapsed features at the species level as well as the summarized abundance of *Bifidobacterium* among pH conditions for each substrate were visualized with bar plots in “ggplot2”.^47^ Differential abundant taxa between different pH programs were found by differential abundance analysis (Deseq2)^48^ with an adjusted p-value lower than 0.05, and differences in the abundance of the top 20 taxa with the lowest adjusted p-values were plotted in a heatmap using the R package “complexheatmap”.^49^ For microbial co-occurrence analysis, species with a mean relative abundance of more than 1.5% were chosen for analysis, and the chord diagram was visualized by R package “circlize” with Spearman’s rank correlation coefficient of more than 0.7. Pearson correlation analysis between metabolite concentrations and the relative abundance of microbial community members was performed with the package “Rhea”,^50^ and R package “corrplot” was used to visualize the correlation coefficients. The minimal prevalence (among the sequenced samples) of one given microbial taxa was set to 30%.

## Funding and Acknowledgement

NMR data were generated through accessing research infrastructure at Aarhus University, including FOODHAY (Food and Health Open Innovation Laboratory, Danish Roadmap for Research Infrastructure). Zhuqing Xie thanks for the finical support from China Scholarship Council (NO. 202006150033) for the PhD study in University of Copenhagen.

## Data Availability Statement

The raw sequences data produced in this study is released through the NCBI Sequence Read Archive under BioProject accession number PRJNA932979. Analytic codes are available upon request.

## Author contribution

DSN formulated the idea of this study and contributed to writing the manuscript; ZQX planned and carried out the experiments, conducted bioinformatics, and wrote the manuscript; WWH assisted ^1^H NMR spectroscopy acquisition and analysis; AG performed Illumina sequencing; ZQX, WWH, AG, HCB, and DSN interpreted data; all authors read, commented on, and approved the final manuscript.

## Abbreviations

BCM: Basal Colon Media
Db-RDA: Distance-based Redundancy Analysis
Deseq2: differential abundance analysis
FOS: Fructooligosaccharides
GM: Gut Microbiota
GOS: Galacto-oligosaccharides
MVDA: Multivariate Data Analysis
OPLS-DA: Orthogonal Projections to Latent Structures Discriminant Analysis
PBS: Phosphate-buffered Saline
SCFA: Short-chain Fatty Acids
TSP: trimethylsilylpropanoic acid
zOTUS: zero radius Operational Taxonomic Units.

