## Supplemental Figures and Tables for "The effect of colonic pH on microbial activity and metabolite production using common prebiotics as substrates: an *in vitro* study"

**Supplementary file**

Table S1. The Q^2^ of OPLS-DA models constructed to discriminate the gut metabolome from different pH groups.

| Comparison | Q^2^ |
| --- | --- |
| Low vs Medium | 0.28 |
| Low vs High | 0.82 |
| Medium vs High | 0.21 |

Q^2^ > 0.5 indicates that the OPLS-DA model is robust.

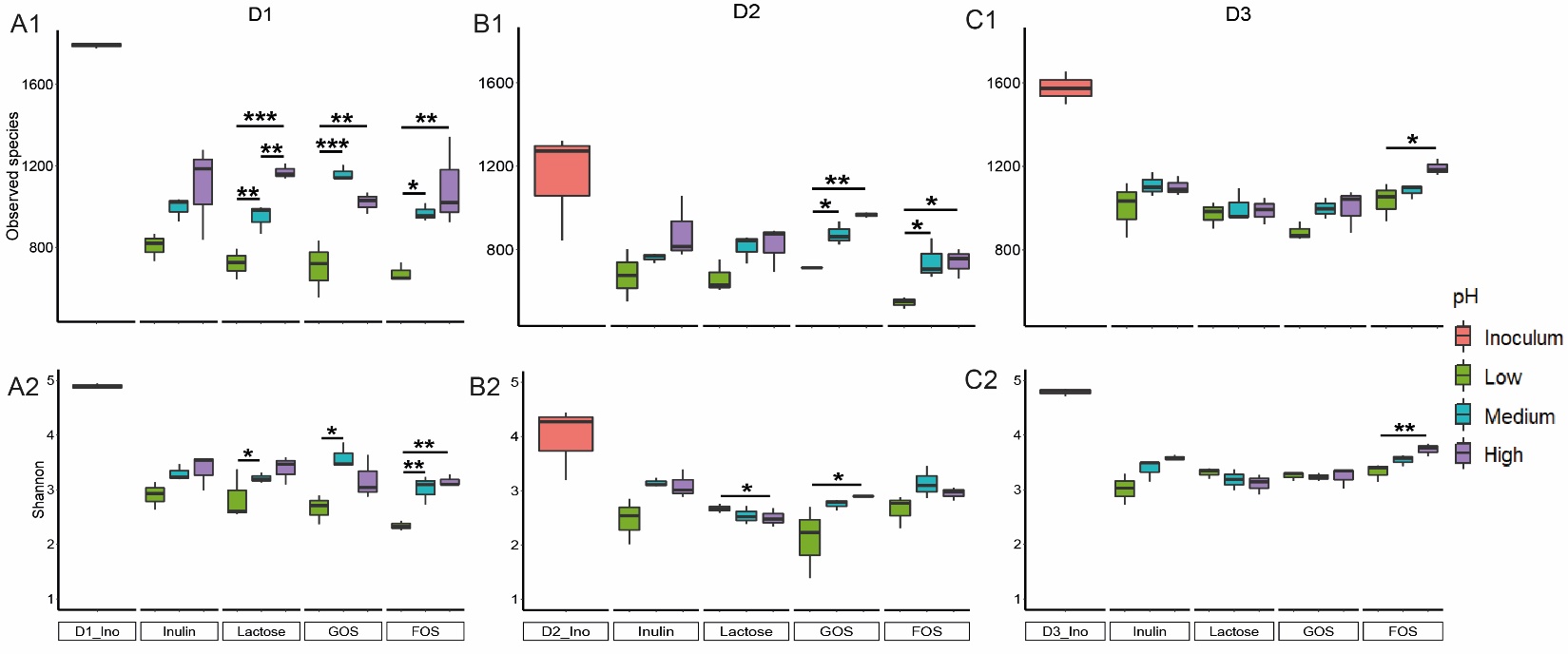

Fig. S1 The influence of colonic pH on individual GM alpha diversity after 24 h of fermentation. A-C represents D1, D2, and D3, and 1-2 represents Observed species and Shannon index. A t-test was applied to determine the influence of pH on alpha diversity for each inoculum with a single substrate. Significant differences between changed colonic pH are labelled with * (*p* < 0.05), ** (*p* < 0.01) and *** (*p* < 0.001), respectively.

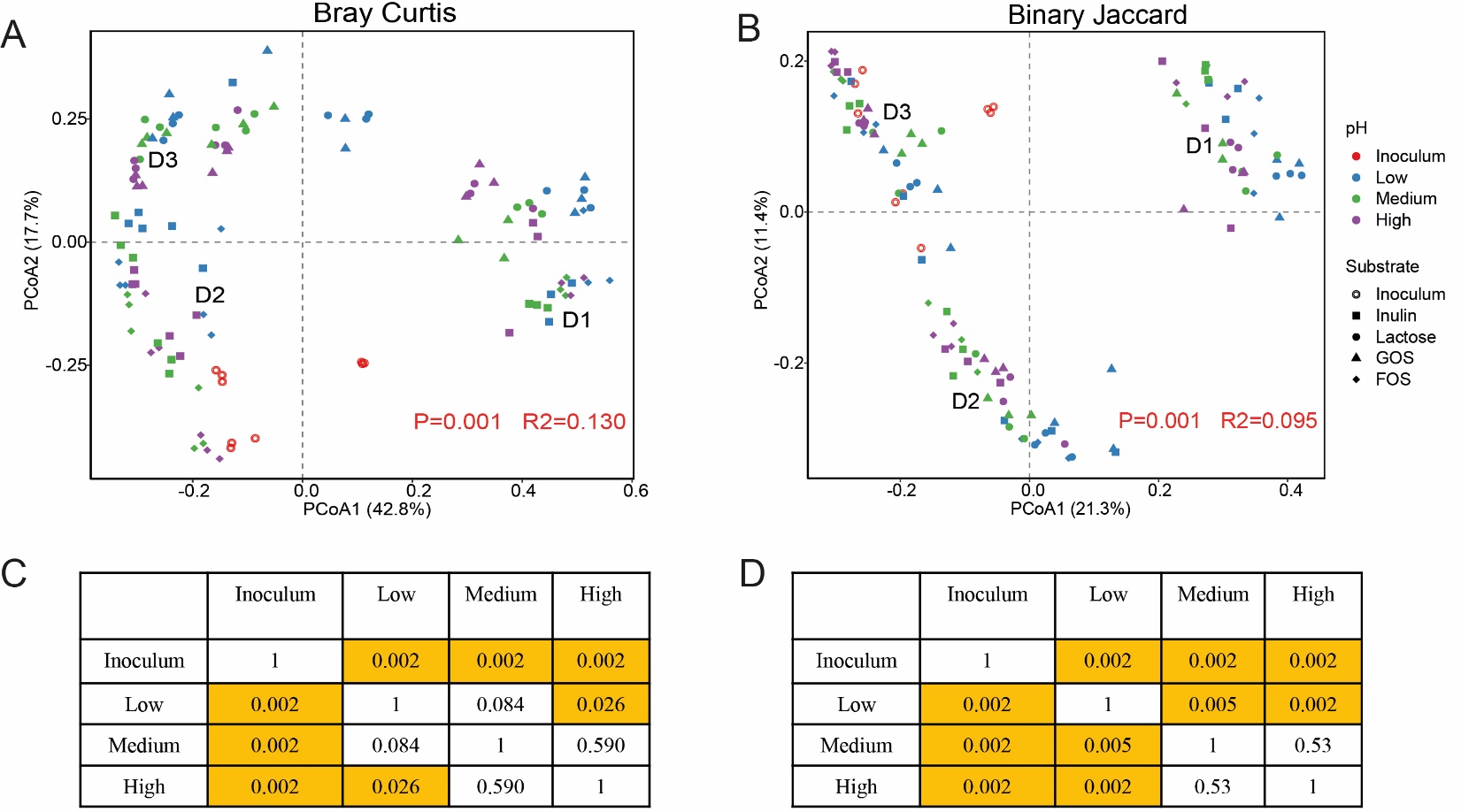

Fig. S2 PCoA and pairwise PERMANOVA tests on Bray Curtis (A and C) and Binary Jaccard metrics (B and D).

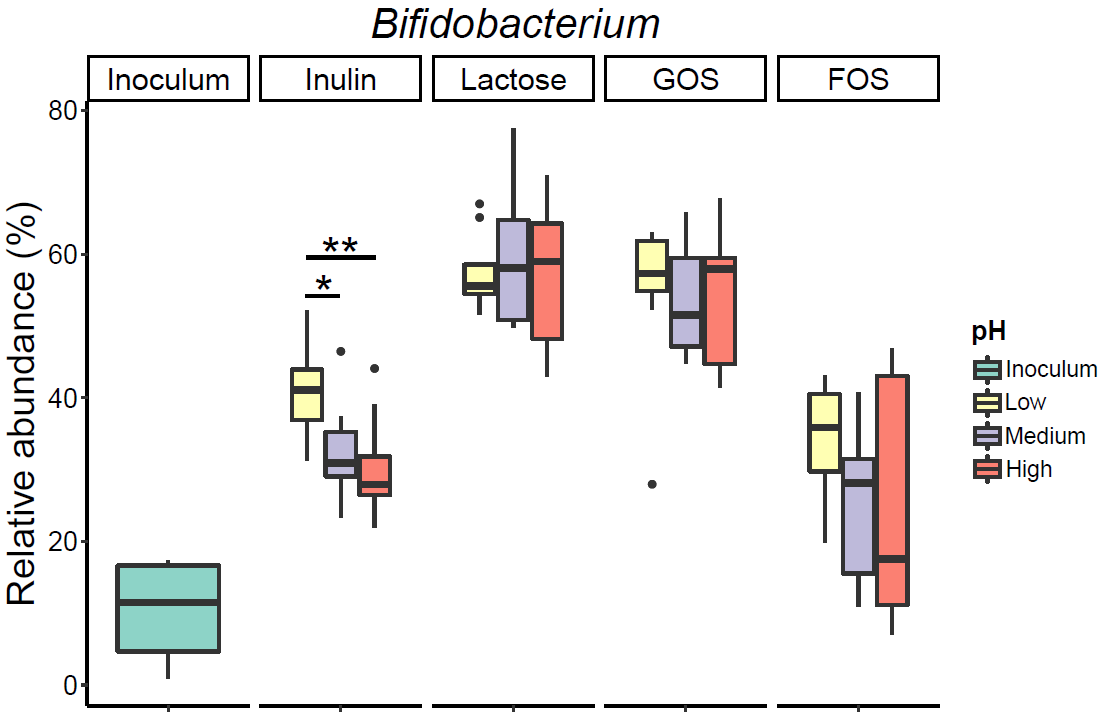

Fig. S3 The influence of colonic pH on *Bifidobacterium* (relative abundance) after 24 h of fermentation using different substrates. A t-test was applied to determine the influence of pH on *Bifidobacterium* with a single substrate. Significant differences between changed colonic pH are labelled with * (*p* < 0.05), ** (*p* < 0.01) and *** (*p* < 0.001), respectively.

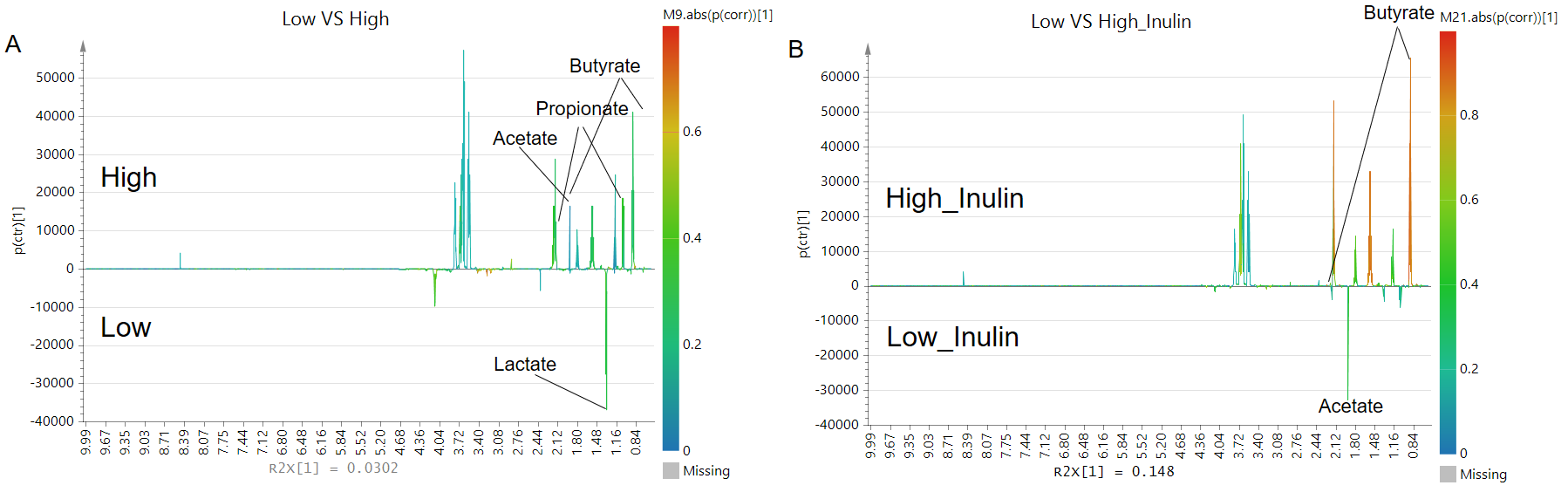

Fig. S4 S-line plot of OPLS-DA visualizing the differences in the NMR metabolite profiles of low and high colonic pH (A, Q^2^ = 0.82) and only inulin (B, Q^2^ = 0.55).

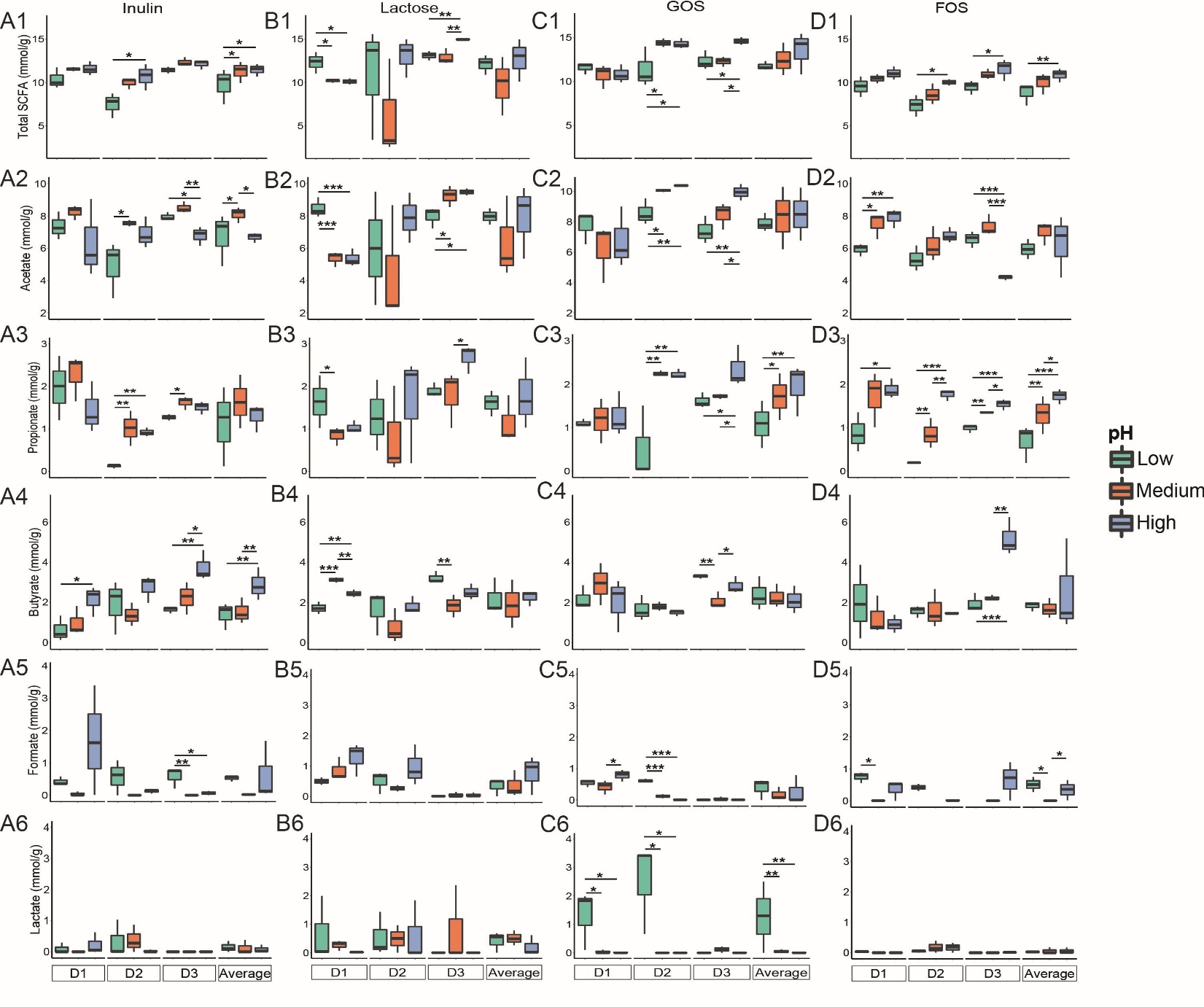

Fig.S5 The influence of colonic pH on individual metabolite production after 24 h of fermentation. A-D represents substrate inulin, lactose, GOS, and FOS, respectively, and 1-6 represents total SCFA, acetate, propionate, butyrate, formate, and lactate production. A t-test was applied to determine the influence of pH on individual metabolite production with a single substrate. Significant differences between changed colonic pH are labelled with * (*p* < 0.05), ** (*p* < 0.01) and *** (*p* < 0.001), respectively.
